# Modular HUWE1 architecture serves as hub for degradation of cell-fate decision factors

**DOI:** 10.1101/2020.08.19.257352

**Authors:** Moritz Hunkeler, Cyrus Y. Jin, Michelle W. Ma, Daan Overwijn, Julie K. Monda, Eric J. Bennett, Eric S. Fischer

## Abstract

HECT ubiquitin ligases play essential roles in metazoan development and physiology. The HECT ligase HUWE1 is central to the cellular stress response by mediating degradation of key death or survival factors including Mcl1, p53, DDIT4, and Myc. As a step toward understanding regulation of HUWE1 engagement with its diverse substrates, we present here the cryo-EM structure of HUWE1, offering a first complete molecular picture of a HECT ubiquitin ligase. The ~4400 amino acid residue polypeptide forms an alpha solenoid-shaped assembly with a central pore decorated with protein interaction modules. This modularity enables HUWE1 to target a wide range of substrates for destruction. The locations of human mutations associated with severe neurodevelopmental disorders link functions of this essential enzyme with its three-dimensional organization.

## Introduction

Ubiquitin E3 ligases confer specificity to the process of ubiquitylation. They fall into to three major classes: RING, RBR, and HECT ligases^1^. HECT ligases (**H**omologous to the **E**6-AP **C**arboxy **T**erminus), which typically have large and complex architectures, are unique in their catalytic mechanism^2^. They are frequently mutated in developmental disorders and are widely implicated in carcinogenesis^3–5^.

The HECT family member HUWE1 (**H**ECT, **U**BA and W**WE** domain containing protein **1**, also known as Mule/ARF-BP1/LASU1/HECTH9) was initially shown to degrade three key regulators of stress and survival: Mcl1, p53, and Myc^6–8^. A growing list of HUWE1 substrates has since been reported^9^, including the stress-responsive regulator of mTORC1 signaling, DDIT4^10,11^, and many DNA damage response factors, such as the BRCA1 tumor suppressor, TopBP1, Cdc6 and CHEK1^12–15^. Loss of HUWE1 sensitizes cells not only to DNA damage, but also to a variety of other stressors, including both oxidative and hypoxic stress^9,16–19^. In addition, HUWE1 has been shown to mediate the destruction of unassembled constituents of multi-protein complexes, contributing to protein quality control^20,21^. Micro-duplications and splice-site or missense variants of the HUWE1 gene are associated with severe forms of intellectual disability (ID) and other developmental defects^3,22^. All of these findings implicate HUWE1 as a central hub of the cellular stress response, growth and apoptosis. To address the question how HUWE1 can engage with such a diverse set of substrates, we determined cryo-electron microscopy (cryo-EM) structures that reveal a ring-shaped E3 ligase architecture and unique modularity.

## Results

### Overall Structure of HUWE1

Before embarking on structural studies, the functional integrity of purified full-length HUWE1 was established by *in vitro* ubiquitylation of the *bona fide* substrates Mcl1 and DDIT4 (Fig. 1a, **Supplementary Fig. 1a-c**). Cryo-EM datasets were collected for BS3-crosslinked (**Supplementary Fig. 1d**) and non-crosslinked samples. Reconstructions were refined to overall resolutions of ~ 3.1 Å and ~ 3.4 Å, respectively (Fig. 1b and **Supplementary Figs. 2-5**). The resulting maps showed nearly indistinguishable structures (**Supplementary Figs. 2, 3a**), and we focus here on the better-defined map from the BS3-crosslinked sample. An atomic model, built manually, was refined against all maps (Fig. 1b-e, **Supplementary Fig. 5e-h, Supplementary Table 1**).

**Fig. 1.**
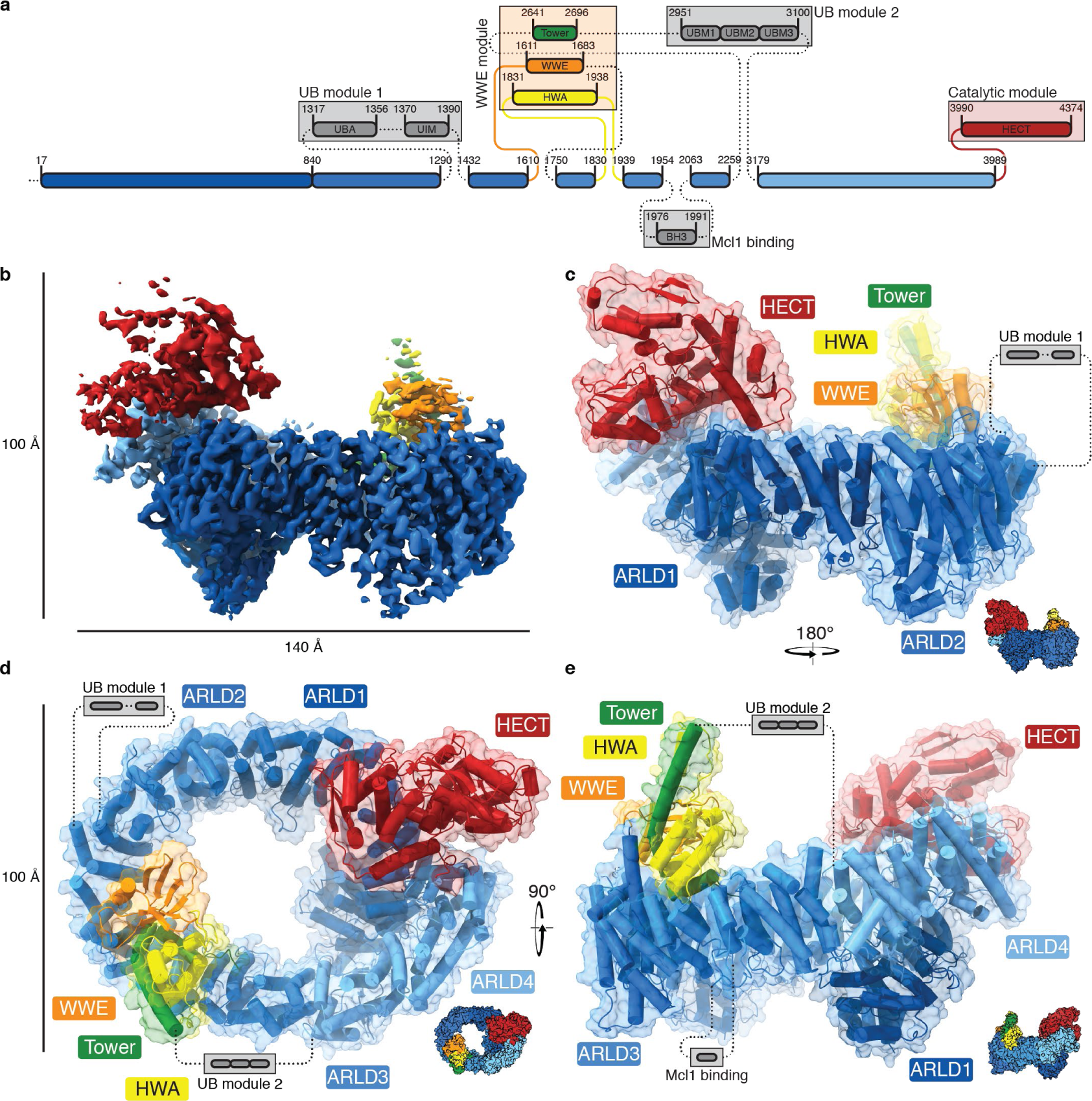
Domain organization and cryo-EM structure of HUWE1. (**a**) Schematic domain overview highlighting the modular nature of HUWE1, with amino residue boundaries indicated. Dotted lines and gray boxes represent unresolved linkers and regions, respectively. (**b**) Cryo-EM density (EMD-22428), sharpened with a B-factor of −20 Å^2^ and shown at a contour level of 0.0159, colored according to color scheme in (a). (**d – e**) Cartoon representation of HUWE1 with transparent surface, domains colored according to scheme in (a). Domains are labeled where applicable, and approximate position and anchoring points for mobile domains are indicated.

The central feature of HUWE1 is a large alpha solenoid structure with dimensions of ~140 Å x ~100 Å x ~100 Å. It is built from four helical **a**rmadillo **r**epeat-**l**ike **d**omains (ARLD1-4), of which two, ARLD1 (amino acid residues (aa) 1-840) and ARLD2 (aa 841-1610), were previously identified based on sequence analysis6. These form, together with ARLD3 (aa 1750-2259) and ARLD4 (aa 3179-3989), a ring-shaped solenoid with an inner circumference of ~250 Å (Fig. 1). This ring is decorated with accessory modules that are inserted in or between the helical repeat domains (Fig. 1a, c-e). The **u**biquitin **b**inding UBA and UIM domains (“UB module 1”, aa 1317-1390) are inserts into ARLD2; they are flexibly tethered to the ring by linkers of 27 and 42 residues. Although not visible in our cryo-EM maps, the UB module 1 is constrained topologically to lie above the plane of the ring (Fig. 1c,d). The WWE domain (aa 1611-1683), which also lies above the plane of the ring, follows ARLD2 (Fig. 1c-e, **Supplementary Fig. 5i**). It interacts with a previously uncharacterized domain (aa 1831-1938, Figs. 1c-e, 2a), inserted into ARLD3, which we termed the “HWA” domain (**H**UWE1 **W**WE module **a**ssociated). The BH3 domain (aa 1976-1991), which lies below the ring, is also an insert into the ARLD3. Like the UB modules, the BH3 domain is not visible in our cryo-EM reconstructions (Figs. 1e, 2a). A largely disordered region spans residues 2259-3179 between ARLD3 and ARLD4, only featuring one folded helix-turn-helix motif (“Tower”, aa 2641-2696, Figs. 1a, 2a) and a UBM domain consisting of three repetitive elements (“UB module 2”, aa 2951-3100, Figs. 1a, 2a). The Tower motif is found tightly sandwiched between the WWE and HWA domains (“WWE module”, Figs. 1c-e, 2a, **Supplementary Fig. 5g**).

**Fig. 2.**
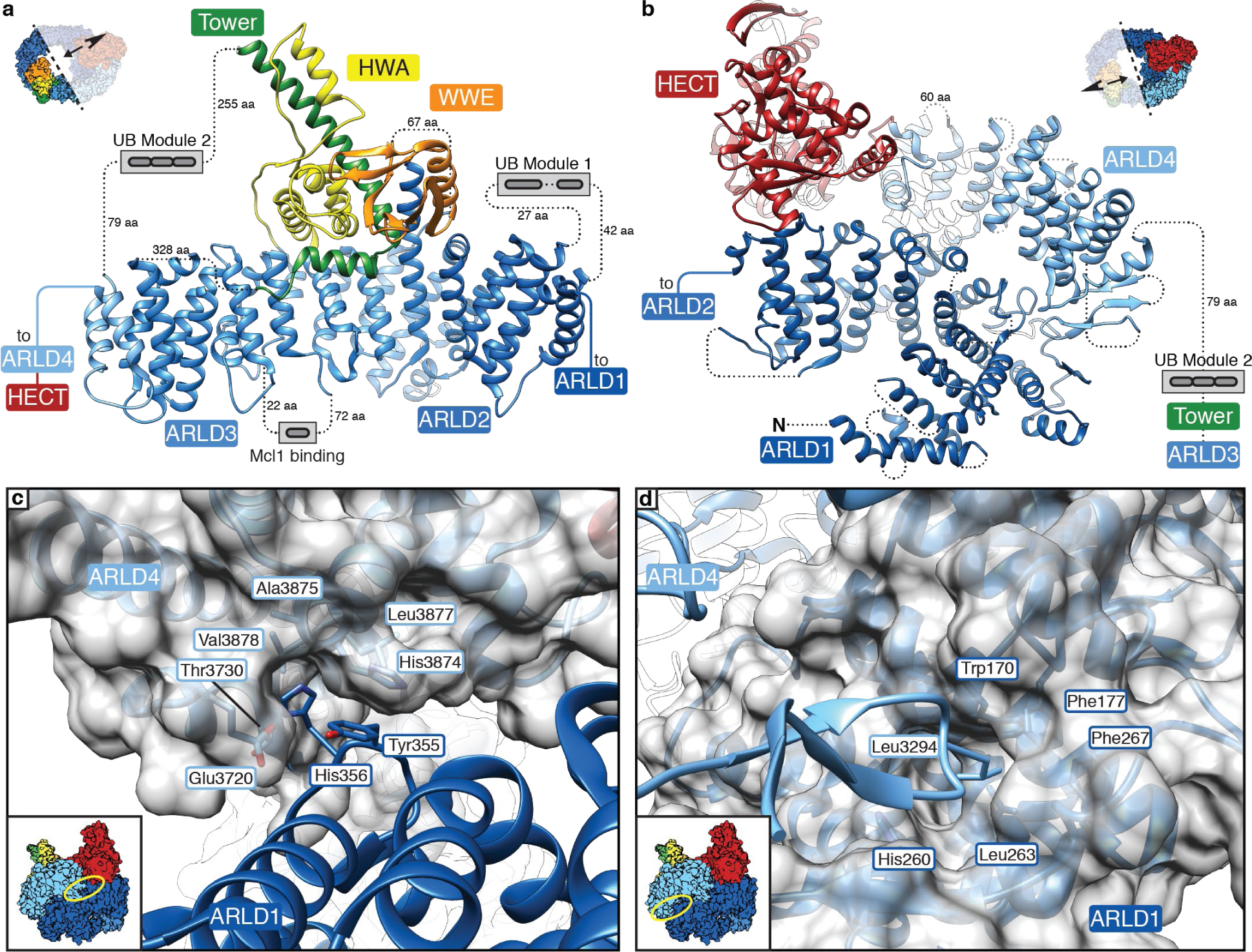
Closed ring architecture of HUWE1. Close-up view of HECT distal (**a**) and HECT proximal (**b**) halves of the HUWE1 structure. All domains are colored and labeled as in Fig. 1a and lengths of unresolved linkers to modules are indicated. (**c, d**) Close-up view of the two key motifs mediating ring closure with transparent surface for ARLD4 and ARLD1 in (c) and in (d), respectively. Key residues forming the interface are labeled.

The HECT domain (“catalytic module”, aa 3990-4374) is above the plane of the ring, exactly opposite the WWE module (Fig. 1c-e). The catalytic mechanism of HECT ubiquitin ligases requires the HECT domain to run through distinct states of a catalytic cycle^2^. In our structure, the HECT domain is found in a resting or inactive conformation, which is compatible with transition into an E2-binding state (**Supplementary Fig. 6a,b**). To move into a ubiquitin transfer-competent state (**Supplementary Fig. 6c**), the HECT domain has to undergo a large conformational rearrangement, which is enabled by the presence of a long (60 aa) flexible linker in ARLD4 (Fig. 2b and **Supplementary Fig. 6d**). This capacity for the domain to move and reorganize is supported by structural variance analysis (**Supplementary Movie 1**).

### HUWE1 architecture forms an activity scaffold

The alpha solenoid architecture is established by a ring-closure interface that comprises residues of the ARLD1 and the ARLD4 domains (Fig. 2b-d), with a total interface area of ~860 Å^2^. Two structural motifs are the principal features of the contact. In one, Tyr355 and His356, in a helix-loop-helix motif in ARLD1, are buried in a largely hydrophobic groove in the ARLD4 domain (Fig. 2c). In the other, an extended loop (aa 3288-3299), with Leu3294 at its tip protrudes from the ARLD4 domain into a hydrophobic pocket in the ARLD1 domain (Fig. 2d). The presence of ‘open’ particles in the non-crosslinked dataset (**Supplementary Fig. 3 and Supplementary Movie 2**) suggested that the ring closure might be subject to regulation, prompting us to investigate whether the presence of a closed ring is critical for activity. Accordant with the functional importance of the closed architecture, mutations in either the helix-loop-helix motif (Y355G/H356G, “YH/GG”) alone or in combination with loop mutations (L3294R and L3294G) (Figs. 2c,d, 3a,b), largely abolished HUWE1 activity. These results suggest that the closed conformation of HUWE1 is essential for activity and indicate that the ring architecture cooperates with distinct binding domains for full activity.

### Protein binding domains control substrate access and function

HUWE1 has at least two domains with substrate-selective features. First, the WWE domain, implicated in directing HUWE1 towards PARylated substrates^23^, facing the center of the solenoid (**Supplementary Fig. 5i**). Second, the BH3 domain, which is known to recruit Mcl1^6^. *In vitro* ubiquitylation assays confirmed that the BH3 domain is necessary for ubiquitylation of Mcl1, but dispensable for ubiquitylation of DDIT4 (Fig. 3c,d). Mutations in the WWE domain did not affect ubiquitylation of Mcl1 nor DDIT4 (Fig. 3c,d). This demonstrates that HUWE1 uses structurally distinct accessory domains that serve as integrated substrate receptors organized on a large scaffold. The limited functional domains, however, are outnumbered by substrates for which there must be additional binding sites. The most likely location for these sites is the ring architecture itself. Armadillo repeats, forming ARLD1-4, are classical protein-protein interaction motifs that bind unfolded or extended peptides on their concave side^24,25^.

**Fig. 3.**
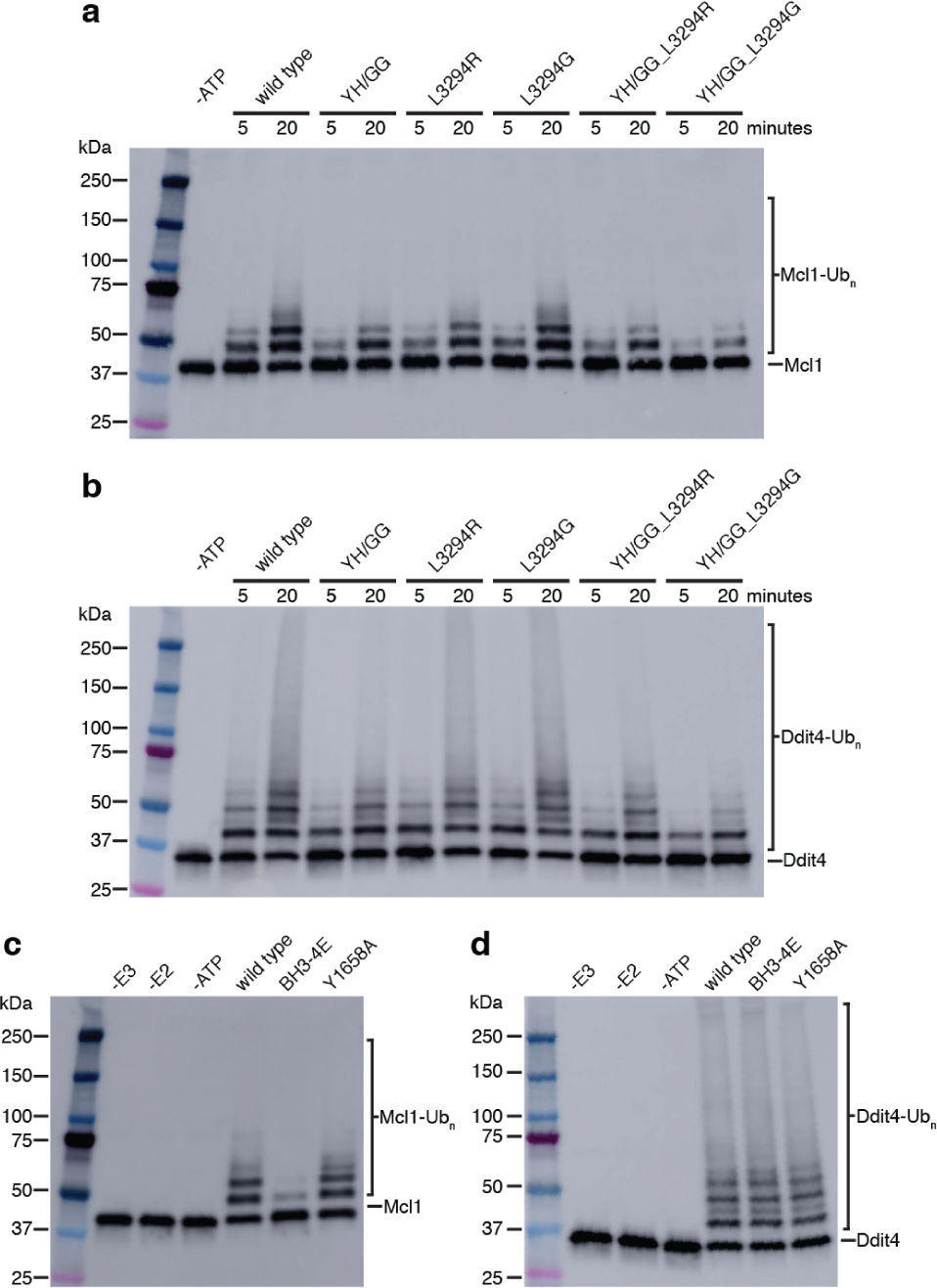
Ring architecture and functional domains are important for activity. (**a, b**) Western blots of *in vitro* ubiquitylation of Mcl1 (a) or DDIT4 (b) by wild type HUWE1, or HUWE1 harboring mutations designed to disrupt the closed ring structure. (**c, d**) *In vitro* ubiquitylation of Mcl1 (c) or DDIT4 (d) by wild type HUWE1 or HUWE1 harboring mutations in the BH3 (V1976E, V1980E, L1983E, M1987E, “BH3-4E”) or WWE (Y1658) domains^6,23^.

### Patient mutations impair ring closure and HUWE1 activity

Mapping patient mutations that lead to ID^3^ onto the structure defines four distinct classes (**Supplementary Fig. 7**). Class I comprises mutations that interfere with ring closure (Figs. 2d, 4a); class II mutations map to the HECT domain. When we introduced these mutations into HUWE1, we found that members of both classes resulted in decreased activity (Fig. 4b,c). Class III mutations cluster in the ARLD1 domain and presumably impair substrate binding. Class IV mutations relate to inactivation of ubiquitin binding activity; when we introduced them into HUWE1, we found them to be either functionally inert or slightly activating (**Supplementary Fig. 8a,b**). Patient mutations in HUWE1 thus have a wide range of functional consequences, from those that slightly increase HUWE1 activity to those that ablate HUWE1 function. The diversity of effects is consistent with the various distinct biological roles of this ligase.

**Fig. 4.**
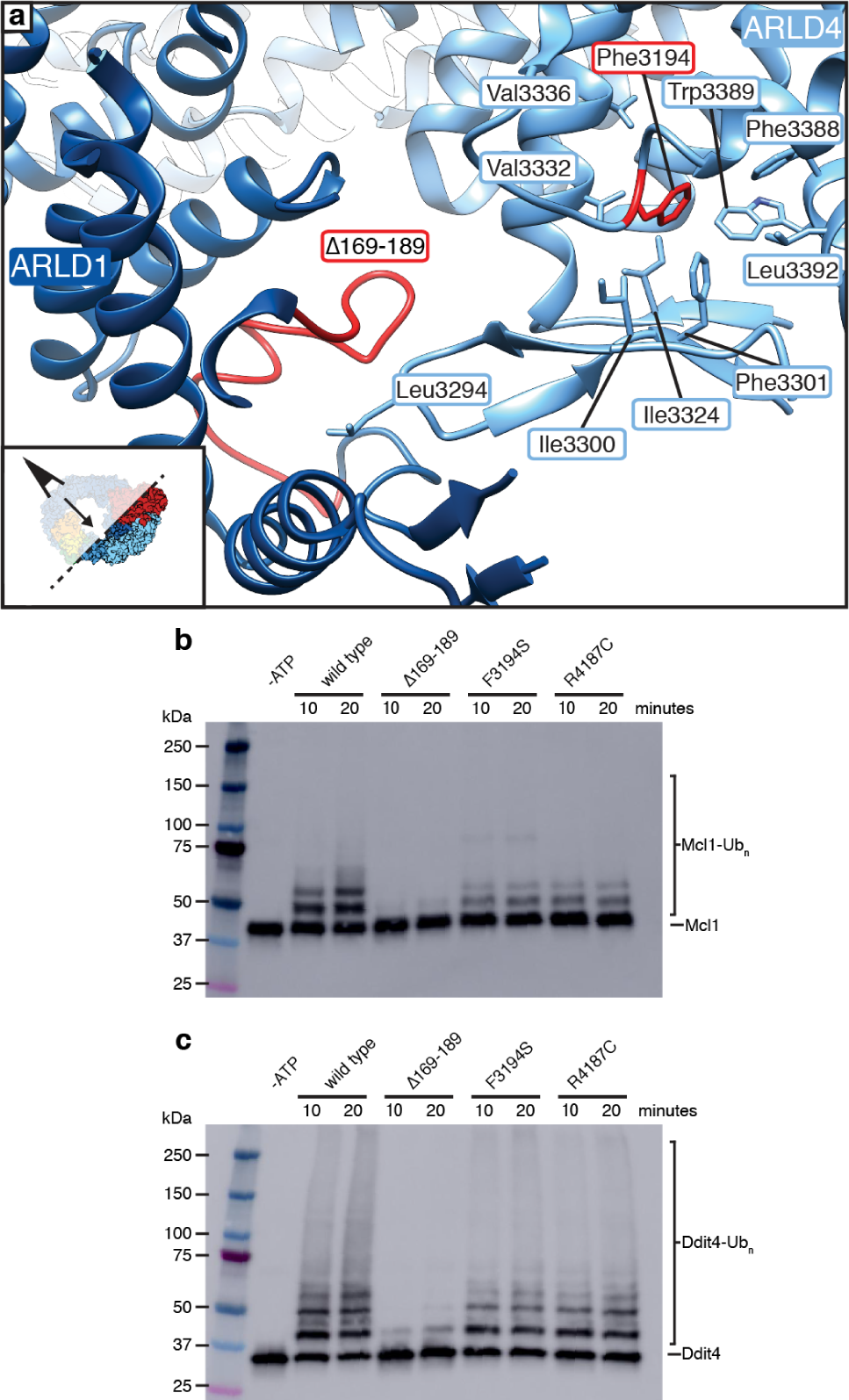
Patient mutations lead to decreased activity. (**a**) Close-up view of the ring closure interface highlighting positions of patient mutations from class I (red). Both the residues stretching aa169-189 as well as Phe3194 are stabilizing the ring closure, by forming part of the interface and by stabilizing the loop conformation, respectively. Hydrophobic residues around Phe3194 are labeled. (**b, c**) *In vitro* ubiquitylation of Mcl1 (b) or DDIT4 (c) by wild type HUWE1 or HUWE1 harboring patient mutations from class I (deletion of aa 169-189 (∆169-189) and F3194S) and class II (R4187C) as indicated.

## Discussion

Our structural and functional characterization of HUWE1, the first of a full-length HECT ligase, reveals a novel, modular architecture for single-chain ubiquitin ligases (Fig. 5a). One conundrum about HUWE1 revolves around its ability to engage with such a large number of substrates. Dedicated binding domains, such as the BH3, the WWE domain and the UB modules, allow for binding of Mcl-1, PARylated and ubiquitylated substrates (Fig. 5b). In our structure we now observe an additional extended protein-protein interaction interface on the inner side of the ring, suggesting that HUWE1 in addition to the protein binding modules uses potentially degenerate peptide binding sites on the helical repeats to access its substrates (Fig. 5c). It is conceivable that multiple binding sites for the same target may exist and that the distinct substrate receptor domains cooperate with the helical repeats, allowing for fine-tuning of substrate recruitment and ubiquitylation activity. Disordered peptide stretches, as well as sites for post-translational modifications (such as ubiquitylation, phosphorylation and PARylation), are observed in most of the reported HUWE1 targets. For example, DDIT4, for which no dedicated substrate receptor domain exists in HUWE1, contains a disordered N-terminal polypeptide stretch (aa 1-84) housing several phosphorylation sites (Fig. 5c). This stretch is dispensable for DDIT4 activity^26^, but its phosphorylation state has been linked to protein stability^27^. In addition to the helical repeats, the large disordered region (aa 2259-3179), which is mostly absent in the closest HUWE1 homolog in yeast^28^ (TOM1), may also contribute to binding towards a specific subset of substrates. The modularity observed in the structural data and evolution of HUWE1^22,28–30^ adapts well to its function as a major cellular rheostat that controls stability of many key regulators of cell growth and death. On the basis of our structure it will now be possible to interrogate the binding modes of a diverse set of substrates in future studies.

**Fig. 5.**
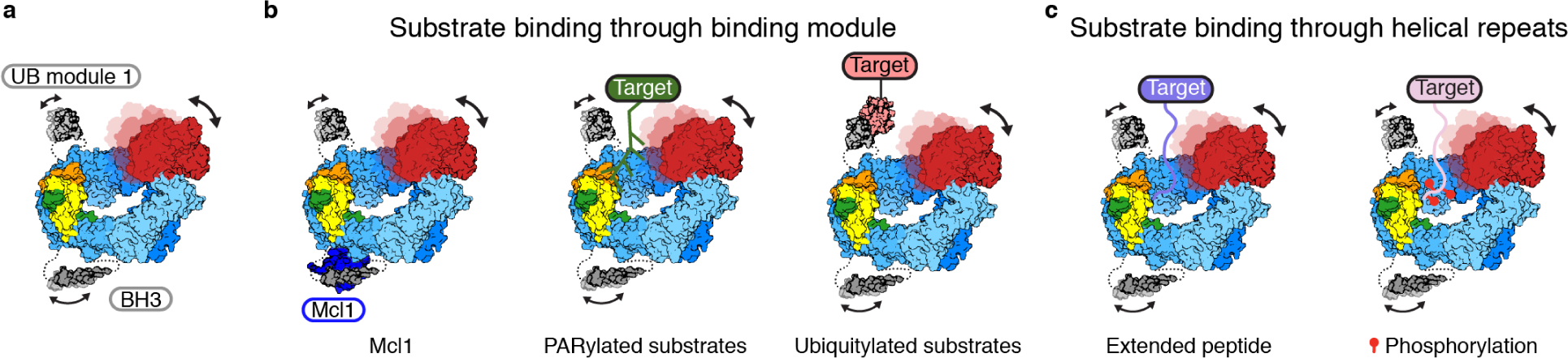
Model for HUWE1 engaging diverse substrates. (**a**) Cartoon representation of HUWE1. UB module 2 and disordered region omitted for clarity. (**b**) Target recruitment through dedicated binding domains involves the BH3 domain (recruiting Mcl-1), the WWE module (recruiting PARylated substrates substrates) and the UB modules (recruiting ubiquitylated substrates). (**c**) The helical repeats may bind to disordered peptide stretches and read out phosphorylation state.

The occurrence of both micro-duplications and missense mutations in patients suggests that both increased (gain of function) and decreased activity of HUWE1 (loss of function) can result in intellectual disability^3,22,31^. Our findings that mutations affecting the architecture or HECT domain are inactivating, while mutations affecting the UB modules are activating or benign, support this hypothesis. None of the patient mutations tested led to a complete loss of activity, in line with the essential role of HUWE1 and the observation that complete loss of HUWE1 is probably not tolerated in humans^32^.

The mechanistic understanding of substrate recruitment and regulation provided here will facilitate the further elucidation of the complex role HUWE1 plays in regulating cellular stress responses^33^. Moreover, as HUWE1 has recently been demonstrated to be a key sensitizer to commonly used anti-cancer agents^16^, our structure may suggest routes to new therapeutics that shift the apoptotic threshold. In particular, it may inform the design of substrate-selective recruiting or inhibiting small molecules.

## Acknowledgments

We thank all the people at the University of Massachusetts Medical School Cryo-EM Core Facility and the Harvard Cryo-EM Center for Structural Biology for their support during grid screening and data collection. We thank everybody involved in the SBGrid consortium for their assistance with high-performance computing. We also thank Michael J. Eck, Nicolas H. Thomä, Nathanael S. Gray, Stephen C. Harrison and Johannes C. Walter for critical feedback on the manuscript.

## Funding

This work was supported by NIH grants NCI R01CA2144608 and R01CA218278 to E.S.F and R01GM127681 to E.J.B.. E.S.F. is a Damon-Runyon Rachleff Investigator (DRR-50-18). J.K.M. was supported by a Postdoctoral Fellowship, PF-19-072-01 – TBE, from the American Cancer Society. M.H. was supported by Swiss National Science Foundation fellowships 174331 and 191053.

## Author contributions

M.H. and E.S.F. designed research plan. M.H. cloned and purified proteins, prepared EM grids, collected data and determined structures. C.J. established, conducted and analyzed activity assays. D.O. assisted with cloning and experiments. M.H., C.J., M.M., D.O., J.M., E.J.B. and E.S.F. contributed to experimental design, analyzed and interpreted data. M.H. prepared figures. E.J.B. and E.S.F. conceived and supervised the study and acquired funding. M.H. and E.S.F. wrote the manuscript with input from all authors. All authors approved the final version of the manuscript.

## Declaration of Interests

E.S.F. is a founder, scientific advisory board, and equity holder in Civetta Therapeutics, Jengu Therapeutics, and Neomorph, Inc. E.S.F. is an equity holder in C4 Therapeutics and a consultant to Novartis, AbbVie, EcoR1 capital, Deerfield, and Astellas. The Fischer lab receives or has received research funding from Novartis, Deerfield and Astellas.

## Data and materials availability

Coordinates have been deposited in the PDB under the accession codes 7JQ9. Cryo-EM maps have been deposited in the EMDB under the accession codes EMD-22427 (BS3-crosslinked, consensus), EMD-22428 (BS3-crosslinked, focus on HECT), EMD-22429 (BS3-crosslinked, focus on WWE), EMD-22430 (BS3-crosslinked, focus on interface), EMD-22431 (non-crosslinked). Uncropped gel and Western blot source data are shown in **Supplementary Figs. 9, 10**.

## Methods

### Cloning, protein expression and purification

A HUWE1 Entry clone (pENTR1A-HUWE1 was a gift from Jean Cook (Addgene plasmid # 37431)) was re-cloned to represent canonical isoform 1 sequence using standard restriction/ligation cloning (amino acids 1-4374, uniprot Q7Z6Z7). HUWE1 mutant sequences were generated by excising pieces from this Entry clone using unique restriction sites, followed by insertion of synthesized double-stranded DNA (gBlocks, IDT) carrying desired mutations using Gibson assembly (New England Biolabs). A modified pDEST plasmid (Thermo Fischer Scientific) was used to transiently express wild-type full-length HUWE1 and all mutants described in this study carrying a single N-terminal FLAG-tag (DYKDDDK) in Expi293 cells (Thermo Fischer Scientific) following the manufacturer’s manual. Cells were harvested 48-60 hours post transfection and lysed by sonication in lysis buffer (50 mM HEPES/KOH pH 7.4, 200 mM NaCl, 2 mM MgCl_2_, 5% glycerol) supplemented with protease inhibitors. The lysate was cleared by ultracentrifugation (45 min, 120,000 g) and incubated with FLAG-antibody-coated beads for 1 hour at 4 °C. Beads were washed with lysis buffer and protein was eluted with 5 column volumes of lysis buffer supplemented with 0.15 mg/ml FLAG peptide. The protein was concentrated using centrifugal concentrators (Amicon, 30 kDa molecular weight cut-off (MWCO)) and polished by size exclusion chromatography (SEC, Superose 6, GE Healthcare) in gel filtration buffer (30 mM HEPES/KOH pH 7.4, 150 mM NaCl, 2 mM TCEP).

Coding sequences of full-length DDIT4 (Harvard plasmid repository, HsCD00327445, amino acid residues 1-232, uniprot Q9NX09) and Mcl-1∆TM (HsCD00004569, amino acid residues 1-327, uniprot Q07820) were subcloned into pAC8-derived vectors^34^ and baculovirus was generated in *Spodoptera frugiperda* Sf9 cells grown in ESF921 medium (Expression systems) following the manufacturer’s manual. Large-scale expression of Strep-tagged proteins for purification was conducted in *Trichoplusia Ni* High Five cells (Thermo Fischer Scientific) in SF-900 medium (Gibco). Cells at a density of 2×10^6^ cells/ml were infected with 1.5% of amplified virus and incubated for 40 hours. After harvest, cells were lysed by sonication in lysis buffer and the lysate was cleared by ultracentrifugation (45 min, 120,000 g). The supernatant was bound to StrepTactin XT High Capacity resin (IBA) in a gravity flow column. The resin was washed with lysis buffer and the proteins were eluted with lysis buffer supplemented with 50 mM biotin. Salt concentration was brought to 50 mM by addition of dilution buffer (50 mM HEPES/KOH pH 7.4, 2 mM TCEP) and proteins were further purified by ion exchange chromatography using a Poros 50HQ (Thermo Fischer Scientific) column, eluting with a linear NaCl-gradient from 50 mM to 750 mM. Protein-containing fractions were combined, concentrated using centrifugal concentrators (Amicon, 10 kDa MWCO), and final polishing was performed by SEC (Superdex200, GE Healthcare) in a buffer containing 30 mM HEPES/KOH pH 7.4, 200 mM NaCl, 2 mM TCEP. During ion exchange chromatography, DDIT4 split into two species (DDIT4 peak1, and DDIT4 peak2), which were both ubiquitylated by HUWE1.

### In vitro ubiquitylation assays

*In vitro* ubiquitylation reactions were carried out in a total volume of 15 μl with UBE1 at 0.2 μM (Boston Biochem), UbcH5b at 0.5 μM (Boston Biochem), wildtype or mutant HUWE1 at 0.21 μM, ubiquitin at 50 μM (Boston Biochem), and Strep-tagged substrate (DDIT4 peak 1, Mcl1) at 2 μM, buffered with 1X E3 Ligase Reaction Buffer (Boston Biochem). Reactions were initiated by addition of Mg-ATP (Boston Biochem) to a final concentration of 10 mM, and were allowed to react for the indicated reaction time at 37 °C. Reaction products were analyzed by SDS-PAGE and subsequent immunoblotting with an anti-Strep antibody HRP conjugate (1:4000, Fischer Scientific). Anti-Strep blots were imaged using an Amersham Imager 600 and Amersham ECL Prime Western Blotting Detection Reagent (GE Life Sciences).

### Cryo-EM sample preparation and data collection

For data set 1 (BS3-crosslinked), HUWE1 eluted from affinity resin was buffer-exchanged using Zeba Spin Desalting Columns (Thermo Fisher Scientific) and incubated with 1.5 mM bis(sulfosuccinimidyl)suberate (BS3, Thermo Fisher Scientific) at room temperature for 20 minutes. Samples were quenched with 100 mM Tris pH 7 before the final SEC step. CHAPSO (Hampton Research) at 0.8 mM was added to HUWE1 at concentrations of 0.9 mg/ml directly before grid preparation. Glow-discharged Quantifoil 1.2/1.3 grids were prepared using a Leica EM-GP, operated at 10 °C and 95% relative humidity. 4 μl samples were applied twice and blotted for 2.25 s each time. Grids were imaged in an FEI Titan Krios equipped with a Gatan Quantum Image filter (20 eV slit width) and a post-GIF Gatan K3 direct electron detector. Images were acquired at 300 kV at a nominal magnification of 105,000 x in counting mode with a pixel size of 0.85 Å/pixel using SerialEM^35^. Three movies (40 frames each) were acquired per hole with four holes per stage position (resulting in 12 image acquisition groups), in a defocus range from −0.8 - −2.5 μm over an exposure time of 2.4 s and a total dose of 45.68 e^−^/Å^2^.

For data set 2 (non-crosslinked), an additional purification step was included between affinity purification and polishing. After elution from FLAG antibody-coated beads, the sample was diluted with dilution buffer to lower the salt concentration (50 mM HEPES/KOH pH 7.4, 2 mM TCEP) and subjected to ion exchange chromatography (Poros 50HQ, Thermo Fischer Scientific), eluted with a linear NaCl-gradient from 50 mM to 750 mM. Peak fractions were pooled, concentrated using centrifugal concentrators (Amicon, 30 kDa MWCO) and polished by SEC. CHAPSO at a concentration of 0.8 mM was added to 0.4 mg/ml HUWE1 directly before grid preparation. 4 μl sample was applied to glow-discharged Quantifoil 1.2/1.3 grids and the grids were vitrified using a Leica EM-GP (3s blotting), operated at 10 °C and 95% relative humidity. Grids were imaged in an FEI Titan Krios equipped with a Gatan Quantum Image filter (20 eV slit width) and a post-GIF Gatan K2 direct electron detector. Images were acquired using SerialEM at 300 kV at a nominal magnification of 130,000 x in super-resolution mode with a pixel size of 0.53 Å per super-resolution pixel at the specimen level. Three movies (35 frames each) were acquired per hole with four holes per stage position (resulting in 12 image acquisition groups), in a defocus range from −1 - −2.3 over an exposure time of 7 s and a total dose of 49.37 e^−^/Å^2^.

### Data processing and model building

Data set 1 (BS3-crosslinked): 10,390 movies were corrected for beam-induced motion using UCSF MotionCor2^36^ (v1.2.1) and contrast transfer function (CTF) was estimated using CTFFIND4.1^37^ (v4.1.10). Poor quality micrographs (CTF resolution estimation >4.9 Å) and micrographs with apparent ice contamination were discarded. 2,110,785 particles were picked from the 9610 remaining micrographs using crYOLO^38^ (v1.2.2) trained on a subset of manually picked particles. Particles were extracted with a box size of 364 pixels (309.5 Å) and down-sampled by Fourier-cropping to a box size of 224 pixels with 1.38 Å/pix in Relion-3.0^39^ (v3.0-beta-2). Particles were cleaned through several rounds of reference-free 2D classification in cryoSPARC^40^ (v2.4.6, unless otherwise noted), and an initial model was calculated from 1,262,934 particles. The particle set (re-extracted with the original pixel size) and resulting initial model were used to further clean up the data set using 3D classification in Relion-3.0. The remaining 762,898 particles went through two rounds of CTF-Refinement and Bayesian polishing implemented in Relion-3.0, globally refining CTF parameters in the first round, followed by CTF-Refinement per image acquisition group in the second round. A consensus refinement yielded a map at 3.3 Å resolution. Masked classification was used to identify subsets of particles with better defined features **(Supplementary Fig. 2)** for the HECT domain (33,078 particles used for reconstruction, 3.8 Å), the WWE/AB/Tower region (85,184 particles, 3.6 Å,) and the N-terminus and interface region (125,477 particles, 3.4 Å,). Available crystal structures of the HECT domain (PDB: 5LP8^41^) and the WWE domain (PDB: 6MIW) could readily be rigid body-fitted into the maps (**Supplementary Fig. 5H**), and all maps were used for manual model building in COOT^42^ (v0.9-pre). Density for the two N-terminal helices (aa 17-27 and aa 31-39) was very weak and did not allow building with high confidence. Two idealized helices based on secondary structure prediction were placed and fitted as rigid bodies into the density. A refined detector pixel size was later determined to be 0.825 Å/pix, so the particle dataset after the first round of Bayesian polishing was imported into Relion-3.1^43^ (v3.1-beta) for reprocessing. In short, beam tilt and anisotropic magnification were refined per image acquisition group. 4^th^ order aberration estimation was used to refine spherical aberration and to estimate the error in the CTF estimation which resulted from using the wrong nominal pixel size in initial processing steps. Refinements resulting in the four maps that were used for model building were repeated, and the resulting maps were re-scaled to a pixel size of 0.825 Å/pix during PostProcess. These final maps (**Supplementary Fig. 2**) were deposited (3.1 Å, EMD-22427; 3.7 Å, EMD-22428; 3.4 Å, EMD-22429; 3.3 Å, EMD-22430). The model was protonated (phenix.reduce) and different parts were individually refined in the maps where they were best resolved (aa 3953-4364 in EMD-22428, aa 1595-2109 and 2512-2696 in EMD-22429, aa 17-376 and aa 3179-4005 in EMD-22430, aa 377-1594 and aa 2109-2259 in EMD-22427). For this, density around the respective part was cut out (using phenix.cut_out_densities^44^ (v1.17.1)) and used for iterative refinement of the atomic model with ISOLDE^45^ (v1.0b4) and phenix.real_space_refine, using adp and rigid body refinement, gradient-driven minimization and simulated annealing. The high-resolution crystal structures of the WWE and the HECT domain as well as the ISOLDE-refined model for the helical repeats were used as target restraints. Finally, the combined model was refined once (3 macro cycles) with target restraints against the map with the highest resolution (EMD-22427). Data collection parameters and final refinement statistics are available as **Supplementary Table 1**.

Data set 2 (non-crosslinked): 9,225 movies were corrected for beam-induced motion and Fourier-cropped by a factor of 2 (resulting in a pixel size of 1.06 Å/pix) using the Relion-3.0 MotionCor implementation, and contrast transfer function (CTF) was estimated using CTFFIND4.1 (v4.1.10). Poor quality micrographs (CTF resolution estimation >5.9 Å) and micrographs with apparent ice contamination were discarded. 2,902,388 particles were picked from the 8404 remaining micrographs by crYOLO (v1.2.2) using the model from data set 1. Particles were extracted with a box size of 288 pixels (305.3 Å) and down-sampled to a box size of 200 pixels with 1.52 Å/pix in Relion-3.0. Particles were cleaned through several rounds of reference-free 2D classification in cryoSPARC, and four initial ab initio models were calculated from 1,356,068 particles. 3 classes represented an open form of HUWE1, and one class represented the closed from observed in data set 1. The increased flexibility of this open form precluded high-quality 3D reconstructions (**Supplementary Movie 2**). Particles in the class representing the closed conformation (~27%) were further cleaned from low-quality particles by calculating 6 additional ab initio models. Good particles (336,853) were re-extracted in Relion-3.0 to the original pixel size, defocus values were estimated per particle in CtfRefine, followed by Bayesian polishing. The resulting particles with enhanced signal were imported back into cryoSPARC for final non-uniform refinement^46^, yielding a map at 3.4 Å (EMD-22431). A correlation of 0.94 between final maps from data set 1 and data set 2 at comparable levels (0.0146 and 0.52, respectively) was determined in Chimera^47^. Data collection parameters are available in **Supplementary Table 1.**

The resolutions of all maps are given based on the Fourier shell correlation (FSC) 0.143 threshold criterion^48,49^. Map and model resolution ranges were judged by local resolution histograms (**Supplementary Figs. 3e. 4c,f,i,l, 7e**). Interface areas were calculated using PDBePisa^50^, and figures of models and EM maps were generated using ChimeraX^51^ and Chimera. 3D structural variability movies^52^ were visualized with Chimera. Structural similarity searches were conducted using PDBeFold^53^, using standard 70% thresholds for both query and target for ARLD1, ARLD3 and ARLD4, while the thresholds were lowered to 40% and 50% for query and target, respectively, for ARLD2. For ARLD1, ARLD2 and ARLD4, armadillo repeat-like proteins populated the top hits. For ARLD3 top hits included ENTH-domain proteins, which are also reminiscent of armadillo repeats^54^. Structural biology applications used in this project were compiled and configured by SBGrid^55^.

## Notes

### Competing Interest Statement

E.S.F. is a founder, scientific advisory board, and equity holder in Civetta Therapeutics, Jengu Therapeutics, and Neomorph. E.S.F. is an equity holder in C4 Therapeutics and a consultant to Novartis, AbbVie, EcoR1 capital, Deerfield, and Astellas. The Fischer lab receives or has received research funding from Novartis, Deerfield and Astellas.

## References

1 Zheng, N. & Shabek, N. Ubiquitin Ligases: Structure, Function, and Regulation. Annu. Rev. Biochem. 86, 129–157 (2017).

2 Lorenz, S. Structural mechanisms of HECT-type ubiquitin ligases. Biol. Chem. 399, 127–145 (2018).

3 Moortgat, S. et al. HUWE1 variants cause dominant X-linked intellectual disability: a clinical study of 21 patients. Eur. J. Hum. Genet. 26, 64–74 (2018).

4 Bernassola, F., Chillemi, G. & Melino, G. HECT-Type E3 Ubiquitin Ligases in Cancer. Trends Biochem. Sci. 44, 1057–1075 (2019).

5 Wang, Y., Argiles-Castillo, D., Kane, E. I., Zhou, A. & Spratt, D. E. HECT E3 ubiquitin ligases - emerging insights into their biological roles and disease relevance. J. Cell Sci. 133 (2020).

6 Zhong, Q., Gao, W., Du, F. & Wang, X. Mule/ARF-BP1, a BH3-only E3 ubiquitin ligase, catalyzes the polyubiquitination of Mcl-1 and regulates apoptosis. Cell 121, 1085–1095 (2005).

7 Adhikary, S. et al. The ubiquitin ligase HectH9 regulates transcriptional activation by Myc and is essential for tumor cell proliferation. Cell 123, 409–421 (2005).

8 Chen, D. et al. ARF-BP1/Mule is a critical mediator of the ARF tumor suppressor. Cell 121, 1071–1083 (2005).

9 Kao, S.-H., Wu, H.-T. & Wu, K.-J. Ubiquitination by HUWE1 in tumorigenesis and beyond. J. Biomed. Sci. 25, 67 (2018).

10 Thompson, J. W. et al. Quantitative Lys-ϵ-Gly-Gly (diGly) Proteomics Coupled with Inducible RNAi Reveals Ubiquitin-mediated Proteolysis of DNA Damage-inducible Transcript 4 (DDIT4) by the E3 Ligase HUWE1. J. Biol. Chem. 289, 28942–28955 (2014).

11 Brugarolas, J. et al. Regulation of mTOR function in response to hypoxia by REDD1 and the TSC1/TSC2 tumor suppressor complex. Genes Dev. 18, 2893–2904 (2004).

12 Wang, X. et al. HUWE1 interacts with BRCA1 and promotes its degradation in the ubiquitin-proteasome pathway. Biochem. Biophys. Res. Commun. 444, 290–295 (2014).

13 Herold, S. et al. Miz1 and HectH9 regulate the stability of the checkpoint protein, TopBP1. EMBO J. 27, 2851–2861 (2008).

14 Cassidy, K. B., Bang, S., Kurokawa, M. & Gerber, S. A. Direct regulation of Chk1 protein stability by E3 ubiquitin ligase HUWE1. FEBS J (2019).

15 Hall, J. R. et al. Cdc6 stability is regulated by the Huwe1 ubiquitin ligase after DNA damage. Mol. Biol. Cell 18, 3340–3350 (2007).

16 Olivieri, M. et al. A genetic map of the response to DNA damage in human cells. bioRxiv, http://www.biorxiv.org/content/10.1101/845446v845441 (2020).

17 Clements, K. E. et al. Identification of regulators of poly-ADP-ribose polymerase (PARP) inhibitor response through complementary CRISPR knockout and activation screens. bioRxiv, http://www.biorxiv.org/content/10.1101/871970v871971 (2019).

18 Bosshard, M. et al. Impaired oxidative stress response characterizes HUWE1-promoted X-linked intellectual disability. Sci. Rep. 7, 15050 (2017).

19 Amici, D. R. et al. Coessential Genetic Networks Reveal the Organization and Constituents of a Dynamic Cellular Stress Response. bioRxiv, http://www.biorxiv.org/content/10.1101/847996v847992 (2019).

20 Xu, Y., Anderson, D. E. & Ye, Y. The HECT domain ubiquitin ligase HUWE1 targets unassembled soluble proteins for degradation. Cell Discov. 2, 16040 (2016).

21 Sung, M. K. et al. A conserved quality-control pathway that mediates degradation of unassembled ribosomal proteins. Elife 5 (2016).

22 Giles, A. C. & Grill, B. Roles of the HUWE1 ubiquitin ligase in nervous system development, function and disease. Neural Dev. 15, 6 (2020).

23 Wang, Z. et al. Recognition of the iso-ADP-ribose moiety in poly(ADP-ribose) by WWE domains suggests a general mechanism for poly(ADP-ribosyl)ation-dependent ubiquitination. Genes Dev. 26, 235–240 (2012).

24 Reichen, C., Hansen, S. & Plückthun, A. Modular peptide binding: from a comparison of natural binders to designed armadillo repeat proteins. J. Struct. Biol. 185, 147–162 (2014).

25 Mönkemeyer, L. et al. Chaperone Function of Hgh1 in the Biogenesis of Eukaryotic Elongation Factor 2. Mol. Cell 74, 88–100.e109 (2019).

26 Vega-Rubin-de-Celis, S. et al. Structural analysis and functional implications of the negative mTORC1 regulator REDD1. Biochemistry 49, 2491–2501 (2010).

27 Katiyar, S. et al. REDD1, an inhibitor of mTOR signalling, is regulated by the CUL4A-DDB1 ubiquitin ligase. EMBO Rep. 10, 866–872 (2009).

28 Marin, I. Origin and evolution of fungal HECT ubiquitin ligases. Sci Rep 8, 6419 (2018).

29 Grau-Bove, X., Sebe-Pedros, A. & Ruiz-Trillo, I. A genomic survey of HECT ubiquitin ligases in eukaryotes reveals independent expansions of the HECT system in several lineages. Genome Biol Evol 5, 833–847 (2013).

30 Marin, I. Animal HECT ubiquitin ligases: evolution and functional implications. BMC Evol. Biol. 10, 56 (2010).

31 Froyen, G. et al. Submicroscopic duplications of the hydroxysteroid dehydrogenase HSD17B10 and the E3 ubiquitin ligase HUWE1 are associated with mental retardation. Am. J. Hum. Genet. 82, 432–443 (2008).

32 Lek, M. et al. Analysis of protein-coding genetic variation in 60,706 humans. Nature 536, 285–291 (2016).

33 Costa-Mattioli, M. & Walter, P. The integrated stress response: From mechanism to disease. Science 368 (2020).

34 Abdulrahman, W. et al. A set of baculovirus transfer vectors for screening of affinity tags and parallel expression strategies. Anal. Biochem. 385, 383–385 (2009).

35 Schorb, M., Haberbosch, I., Hagen, W. J. H., Schwab, Y. & Mastronarde, D. N. Software tools for automated transmission electron microscopy. Nat. Methods 16, 471–477 (2019).

36 Zheng, S. Q. et al. MotionCor2: anisotropic correction of beam-induced motion for improved cryo-electron microscopy. Nat. Methods 14, 331–332 (2017).

37 Rohou, A. & Grigorieff, N. CTFFIND4: Fast and accurate defocus estimation from electron micrographs. J. Struct. Biol. 192, 216–221 (2015).

38 Wagner, T. et al. SPHIRE-crYOLO is a fast and accurate fully automated particle picker for cryo-EM. Commun. Biol. 2, 218 (2019).

39 Zivanov, J. et al. New tools for automated high-resolution cryo-EM structure determination in RELION-3. Elife 7 (2018).

40 Punjani, A., Rubinstein, J. L., Fleet, D. J. & Brubaker, M. A. cryoSPARC: algorithms for rapid unsupervised cryo-EM structure determination. Nat. Methods 14, 290–296 (2017).

41 Sander, B., Xu, W., Eilers, M., Popov, N. & Lorenz, S. A conformational switch regulates the ubiquitin ligase HUWE1. eLife 6, 36 (2017).

42 Emsley, P., Lohkamp, B., Scott, W. G. & Cowtan, K. Features and development of Coot. Acta Crystallogr. D Biol. Crystallogr. 66, 486–501 (2010).

43 Zivanov, J., Nakane, T. & Scheres, S. H. W. Estimation of high-order aberrations and anisotropic magnification from cryo-EM data sets in RELION-3.1. IUCrJ 7, 253–267 (2020).

44 Afonine, P. V. et al. Real-space refinement in PHENIX for cryo-EM and crystallography. Acta Crystallogr. D Struct. Biol. 74, 531–544 (2018).

45 Croll, T. I. ISOLDE: a physically realistic environment for model building into low-resolution electron-density maps. Acta Crystallogr. D Struct. Biol. 74, 519–530 (2018).

46 Punjani, A., Zhang, H. & Fleet, D. J. Non-uniform refinement: Adaptive regularization improves single particle cryo-EM reconstruction. bioRxiv, doi: https://doi.org/10.1101/2019.1112.1115.877092 (2019).

47 Pettersen, E. F. et al. UCSF Chimera--a visualization system for exploratory research and analysis. J. Comput. Chem. 25, 1605–1612 (2004).

48 Scheres, S. H. & Chen, S. Prevention of overfitting in cryo-EM structure determination. Nat Methods 9, 853–854 (2012).

49 Rosenthal, P. B. & Henderson, R. Optimal determination of particle orientation, absolute hand, and contrast loss in single-particle electron cryomicroscopy. J. Mol. Biol. 333, 721–745 (2003).

50 Krissinel, E. & Henrick, K. Inference of macromolecular assemblies from crystalline state. J. Mol. Biol. 372, 774–797 (2007).

51 Goddard, T. D. et al. UCSF ChimeraX: Meeting modern challenges in visualization and analysis. Protein Sci. 27, 14–25 (2018).

52 Punjani, A. & Fleet, D. J. 3D Variability Analysis: Directly resolving continuous flexibility and discrete heterogeneity from single particle cryo-EM images. bioRxiv, http://www.biorxiv.org/content/10.1101/2020.1104.1108.032466v032461 (2020).

53 Krissinel, E. & Henrick, K. Secondary-structure matching (SSM), a new tool for fast protein structure alignment in three dimensions. Acta Crystallogr. D Biol. Crystallogr. 60, 2256–2268 (2004).

54 Legendre-Guillemin, V., Wasiak, S., Hussain, N. K., Angers, A. & McPherson, P. S. ENTH/ANTH proteins and clathrin-mediated membrane budding. J. Cell Sci. 117, 9–18 (2004).

55 Morin, A. et al. Collaboration gets the most out of software. Elife 2, e01456 (2013).

