## Supplementary Information for "Modular HUWE1 architecture serves as hub for degradation of cell-fate decision factors"

Supplementary Figures 1-10

Supplementary Table 1

Legends for Supplementary Movies 1-2

References for Supplementary Material

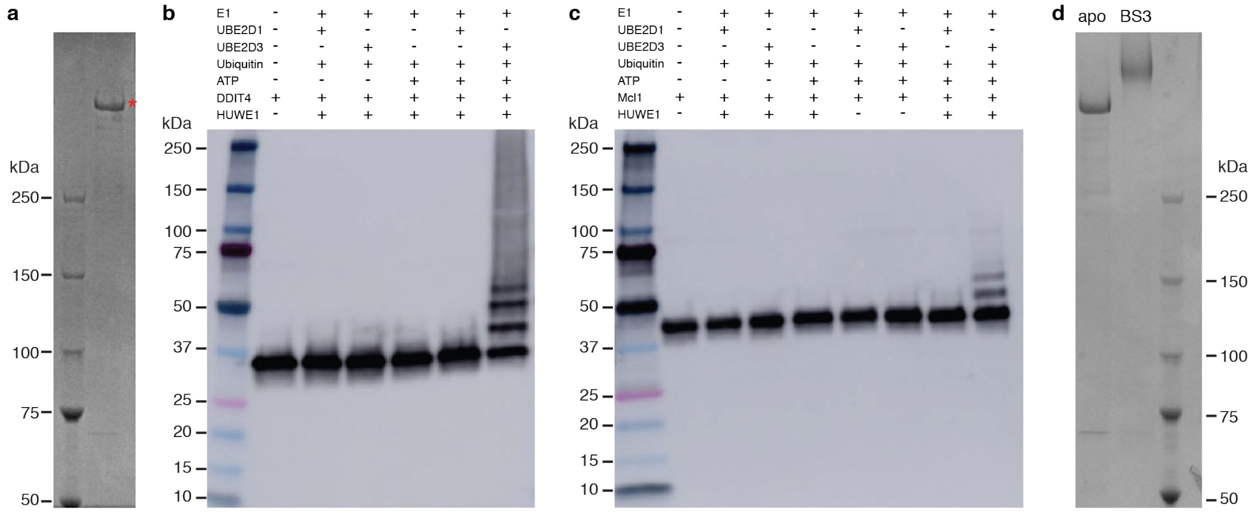

**Supplementary Fig. 1. HUWE1 purification, *in vitro* ubiquitylation, and crosslinking.** (**a**) SDS-PAGE analysis of purified HUWE1 (marked with asterisk). (**b**) *In vitro* ubiquitylation reaction establishing DDIT4 as substrate. Two different E2 were tested. (**c**) *In vitro* ubiquitylation reaction establishing Mcl1 as substrate. Two different E2 were tested. (**d**) SDS-PAGE analysis of non-crosslinked (apo) and BS3-crosslinked HUWE1 illustrating complete crosslinking

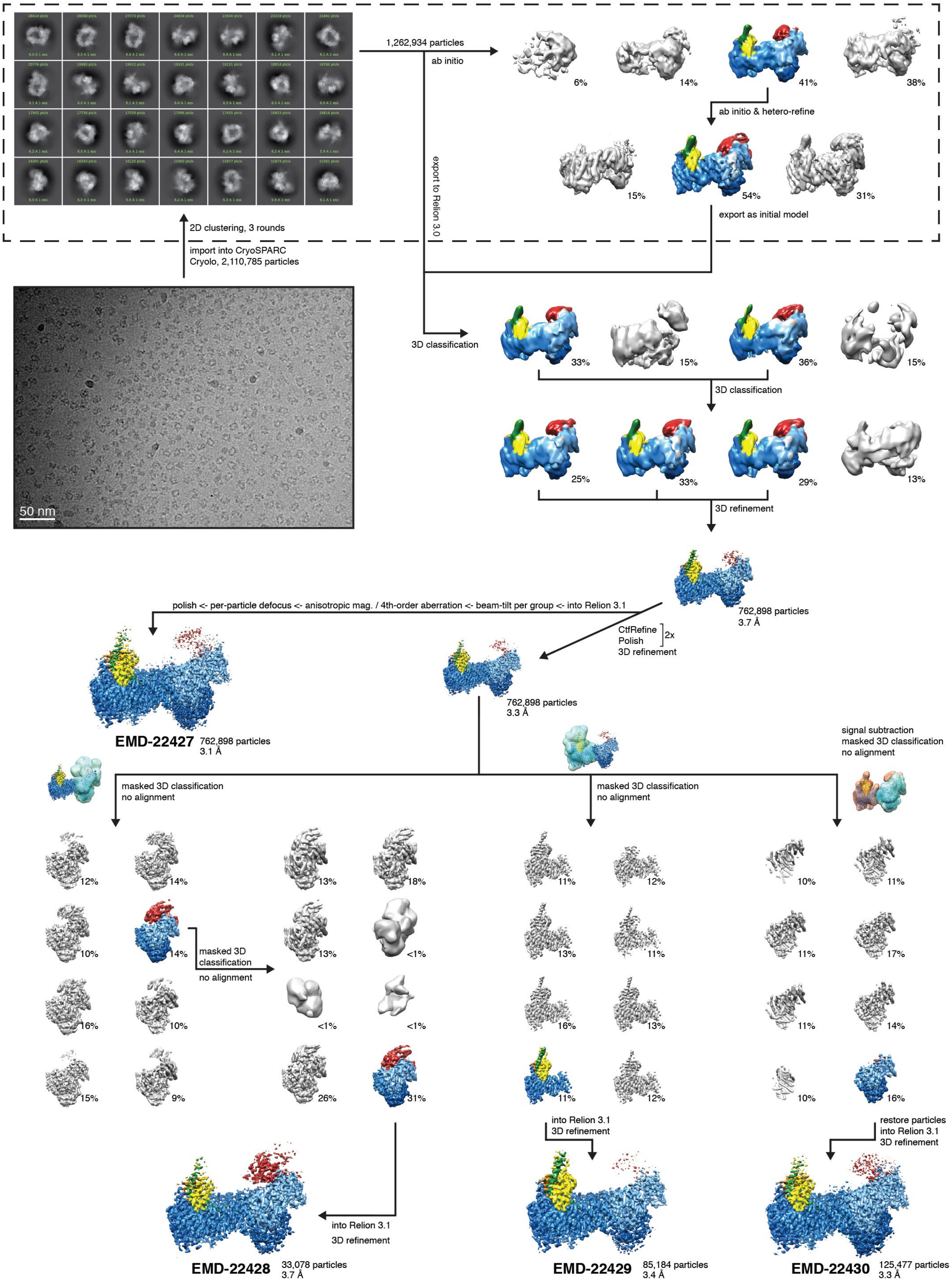
1,262,934 particles
ab initio
6%
14%
41%
38%
ab initio & hetero-refine
15%
54%
31%
export as initial model
export to Relion 3.0
2D clustering, 3 rounds
import into CryoSPARC
Cryolo, 2,110,785 particles
50 nm
3D classification
33%
15%
36%
15%
3D classification
25%
33%
29%
13%
3D refinement
762,898 particles
3.7 Å
polish <- per-particle defocus <- anisotropic mag. / 4th-order aberration <- beam-tilt per group <- into Relion 3.1
CtfRefine
Polish
3D refinement
2x
762,898 particles
3.3 Å
EMD-22427
762,898 particles
3.1 Å
signal subtraction
masked 3D classification
no alignment
masked 3D classification
no alignment
masked 3D classification
no alignment
12%
14%
13%
18%
10%
14%
masked 3D
classification
no alignment
13%
<1%
16%
10%
<1%
<1%
15%
9%
26%
31%
11%
12%
13%
11%
16%
13%
11%
12%
10%
11%
11%
17%
11%
14%
10%
16%
into Relion 3.1
3D refinement
restore particles
into Relion 3.1
3D refinement
into Relion 3.1
3D refinement
EMD-22428
33,078 particles
3.7 Å
EMD-22429
85,184 particles
3.4 Å
EMD-22430
125,477 particles
3.3 Å

**Supplementary Fig. 2. Cryo-EM processing of BS3-crosslinked HUWE1.** Overview of cryo-EM processing workflow, from representative raw micrograph (low-pass filtered to 10 Å) to final deposited maps. Steps enclosed in dashed box were conducted in cryoSPARC, all following steps in Relion3.0 or Relion3.1, as indicated. Particles from 3D classes depicted in color were taken into the next round of processing, and all resolutions indicated are after final PostProcessing. Final sharpening B-factors of deposited maps depicted here are -78, -63, -60 and -75 $A^2$, respectively. Final maps are shown at a contour level of 0.022.

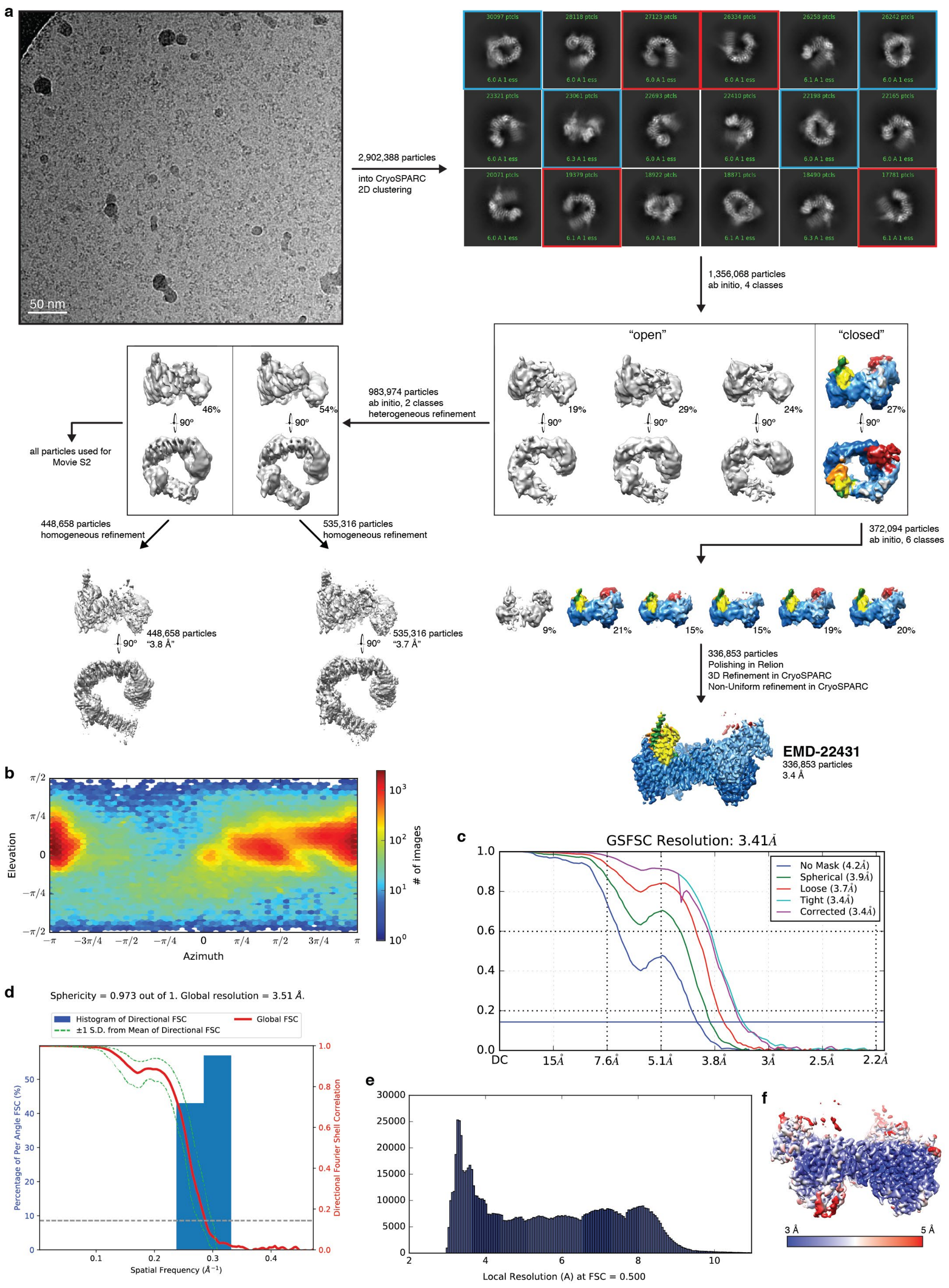
a
50 nm
2,902,388 particles
into CryoSPARC
2D clustering
1,356,068 particles
ab initio, 4 classes
"open"
"closed"
19%
29%
24%
27%
90º
983,974 particles
ab initio, 2 classes
heterogeneous refinement
46%
54%
all particles used for
Movie S2
448,658 particles
homogeneous refinement
535,316 particles
homogeneous refinement
448,658 particles
"3.8 Å"
535,316 particles
"3.7 Å"
372,094 particles
ab initio, 6 classes
9%
21%
15%
15%
19%
20%
336,853 particles
Polishing in Relion
3D Refinement in CryoSPARC
Non-Uniform refinement in CryoSPARC
EMD-22431
336,853 particles
3.4 Å
b
Elevation
Azimuth
### of images
c
GSFSC Resolution: 3.41Å
No Mask (4.2Å)
Spherical (3.9Å)
Loose (3.7Å)
Tight (3.4Å)
Corrected (3.4Å)
DC
15Å
7.6Å
5.1Å
3.8Å
3Å
2.5Å
2.2Å
d
Sphericity = 0.973 out of 1. Global resolution = 3.51 Å.
Histogram of Directional FSC
±1 S.D. from Mean of Directional FSC
Global FSC
Percentage of Per Angle FSC (%)
Directional Fourier Shell Correlation
Spatial Frequency (Å$^{-1}$)
e
Local Resolution (A) at FSC = 0.500
f
3 Å
5 Å

**Supplementary Fig 3. Cryo-EM processing of non-crosslinked HUWE1.** (**a**) Overview of cryo-EM processing workflow, from representative raw micrograph (low-pass filtered to 10 Å) to final deposited map. All steps except polishing were conducted in cryoSPARC. 2D classes show closed (marked with blue boxes) and open (marked with red boxes) HUWE1 particles. Closed particles allowed a reconstruction (sharpening B-factor -116 $A^2$, contour level 0.48) at 3.4 Å, while open particles only led to highly anisotropic reconstructions with heavily over-estimated resolution. Further classification of the particles into large number of classes did not lead to better reconstructions, most probably owing to very high degree of continuous flexibility (**Supplementary Movie 2**). It cannot be excluded that the open form represents an artefact from grid preparation. (**b**) Angular distribution plot for final reconstruction. (**c**) FSC plot for final reconstruction. (**d**) 3DFSC[56] plot and directional resolution histogram for final reconstruction, illustrating high resolution isotropy. (**e**) Local resolution histogram for final reconstruction. (**f**) Final reconstruction colored according to local resolution.

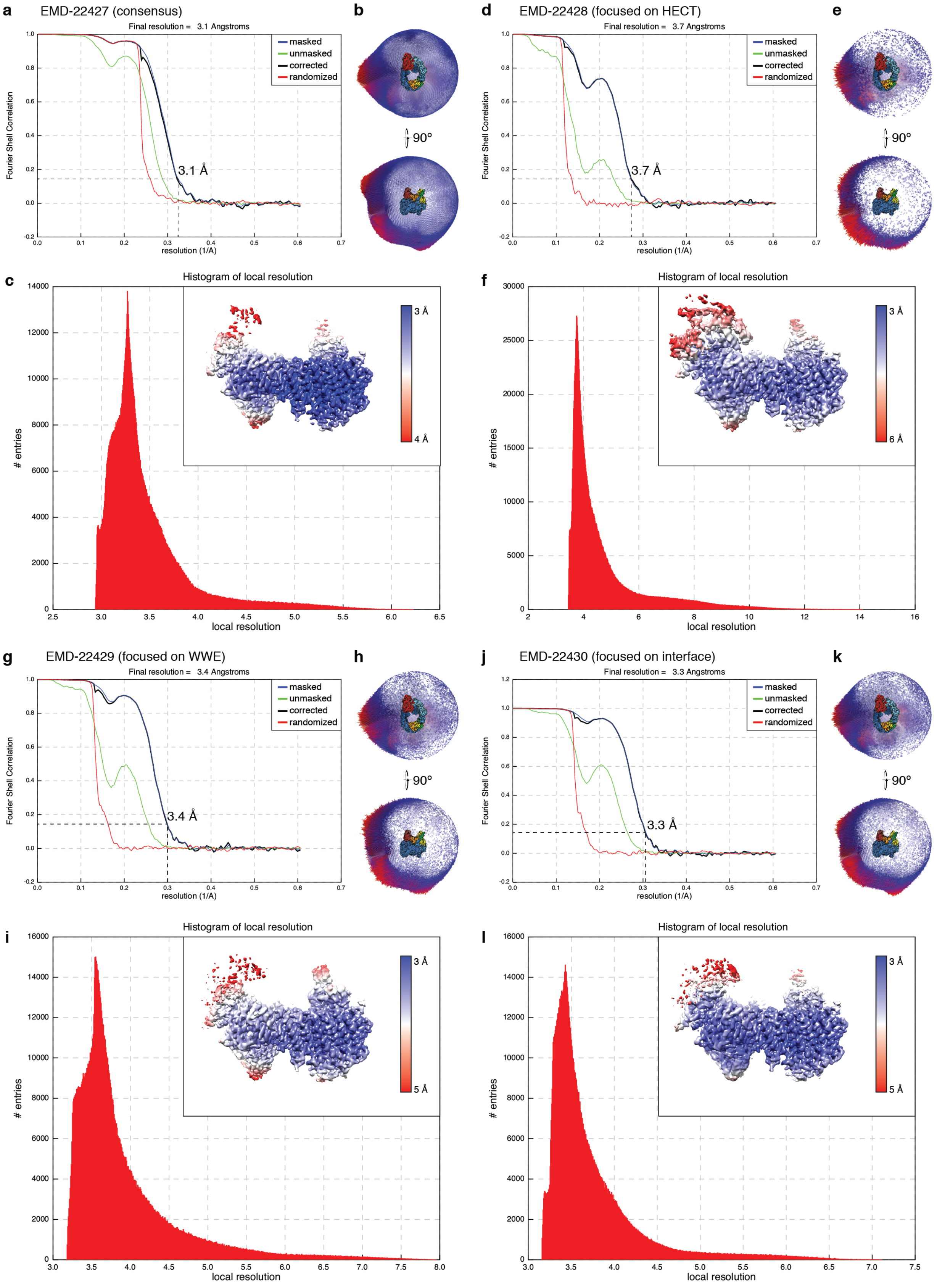
a
EMD-22427 (consensus)
Final resolution = 3.1 Angstroms
masked
unmasked
corrected
randomized
Fourier Shell Correlation
resolution (1/A)
3.1 Å
b
90°
c
Histogram of local resolution
### entries
local resolution
3 Å
4 Å
d
EMD-22428 (focused on HECT)
Final resolution = 3.7 Angstroms
3.7 Å
e
f
6 Å
g
EMD-22429 (focused on WWE)
Final resolution = 3.4 Angstroms
3.4 Å
h
i
5 Å
j
EMD-22430 (focused on interface)
Final resolution = 3.3 Angstroms
3.3 Å
k
l

**Supplementary Fig. 4. Data quality and local resolution for BS3-crosslinked HUWE1.** (**a,d,g,j**), FSC plots for the four deposited maps (EMD-22427(a), EMD-22428 (d), EMD-22429 (g), EMD-22430(j)) of BS3-crosslinked HUWE1, with the corresponding angular distributions (**b, e, h, k**). (**c, f, i, l**) local resolution histograms and, in insets, local resolution mapped onto the cryo-EM density (all maps sharpened with a B-factor of -20 Å$^2$ and shown at contour level of 0.0111).

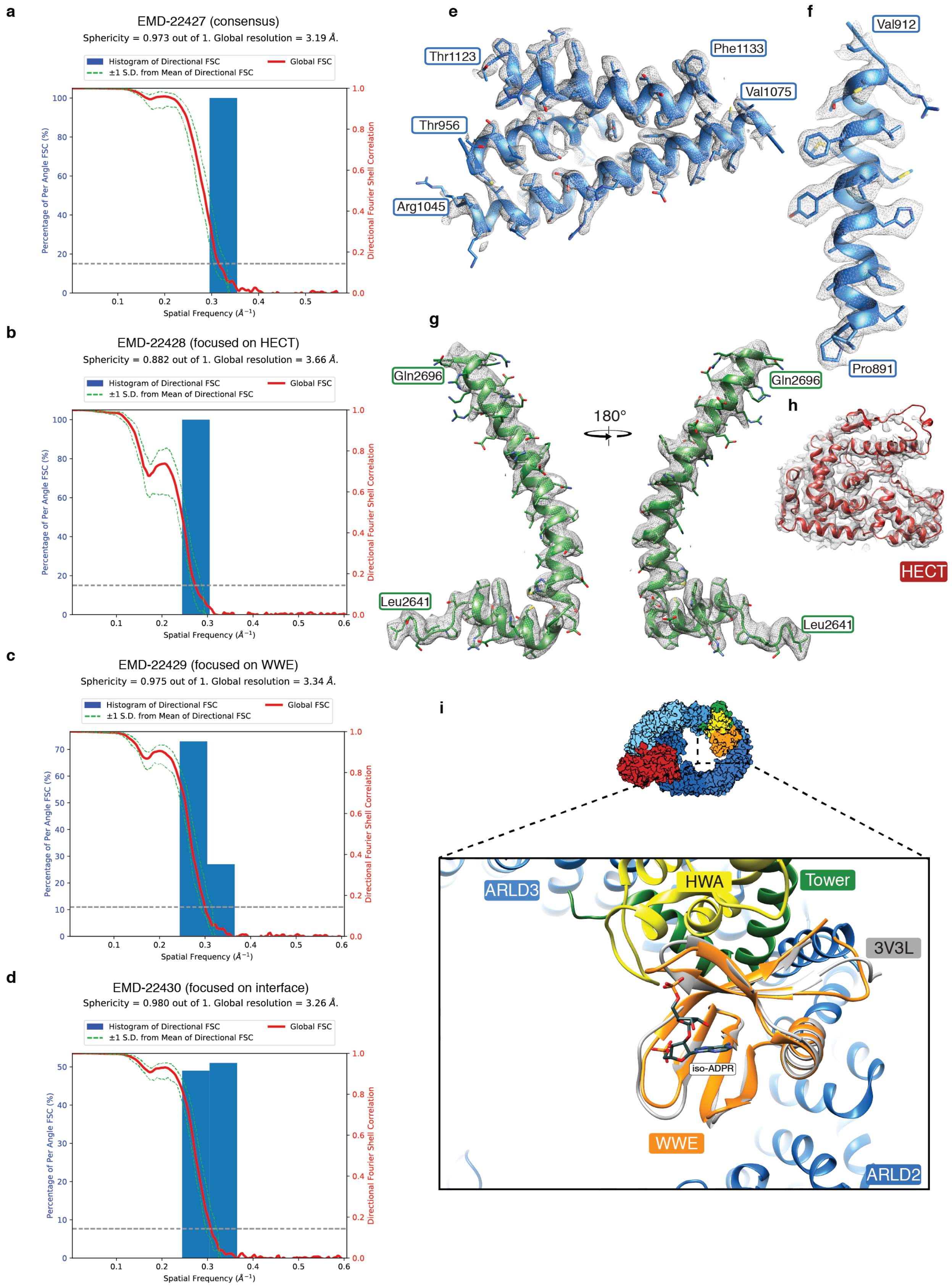
a
EMD-22427 (consensus)
Sphericity = 0.973 out of 1. Global resolution = 3.19 Å.
Histogram of Directional FSC
Global FSC
±1 S.D. from Mean of Directional FSC
Percentage of Per Angle FSC (%)
Directional Fourier Shell Correlation
Spatial Frequency (Å⁻¹)
b
EMD-22428 (focused on HECT)
Sphericity = 0.882 out of 1. Global resolution = 3.66 Å.
c
EMD-22429 (focused on WWE)
Sphericity = 0.975 out of 1. Global resolution = 3.34 Å.
d
EMD-22430 (focused on interface)
Sphericity = 0.980 out of 1. Global resolution = 3.26 Å.
e
Thr1123
Phe1133
Val1075
Thr956
Arg1045
f
Val912
Pro891
g
Gln2696
180°
Gln2696
Leu2641
Leu2641
h
HECT
i
ARLD3
HWA
Tower
3V3L
iso-ADPR
WWE
ARLD2

**Supplementary Fig. 5. Directional FSC, model-density fit and orientation of WWE.** (**a-d**) 3DFSC plot and directional resolution histogram for the four deposited maps of BS3-crosslinked HUWE1, illustrating isotropic resolution distribution in the final reconstructions. (**e,f**), Two representative pieces of EM density with corresponding model of ARLD2 (boundaries indicated) illustrating the map quality in the helical repeats (sharpening B-factor of -78 $Å^2$). (**g**) Density and model of Tower motif (sharpening B-factor of -20 $Å^2$). (**h**) Density and docked HECT domain (sharpening B-factor of -20 $Å^2$). Maps in panels e-h shown at contour level 0.0267. (**i**) Overlay of iso-ADPR-bound WWE domain of RNF146 (PDB: 3V3L[23], gray) with the WWE domain of HUWE1 (orange) in the context of the full-length protein, illustrating that the iso-ADPR binding site is facing the center of the ring architecture.

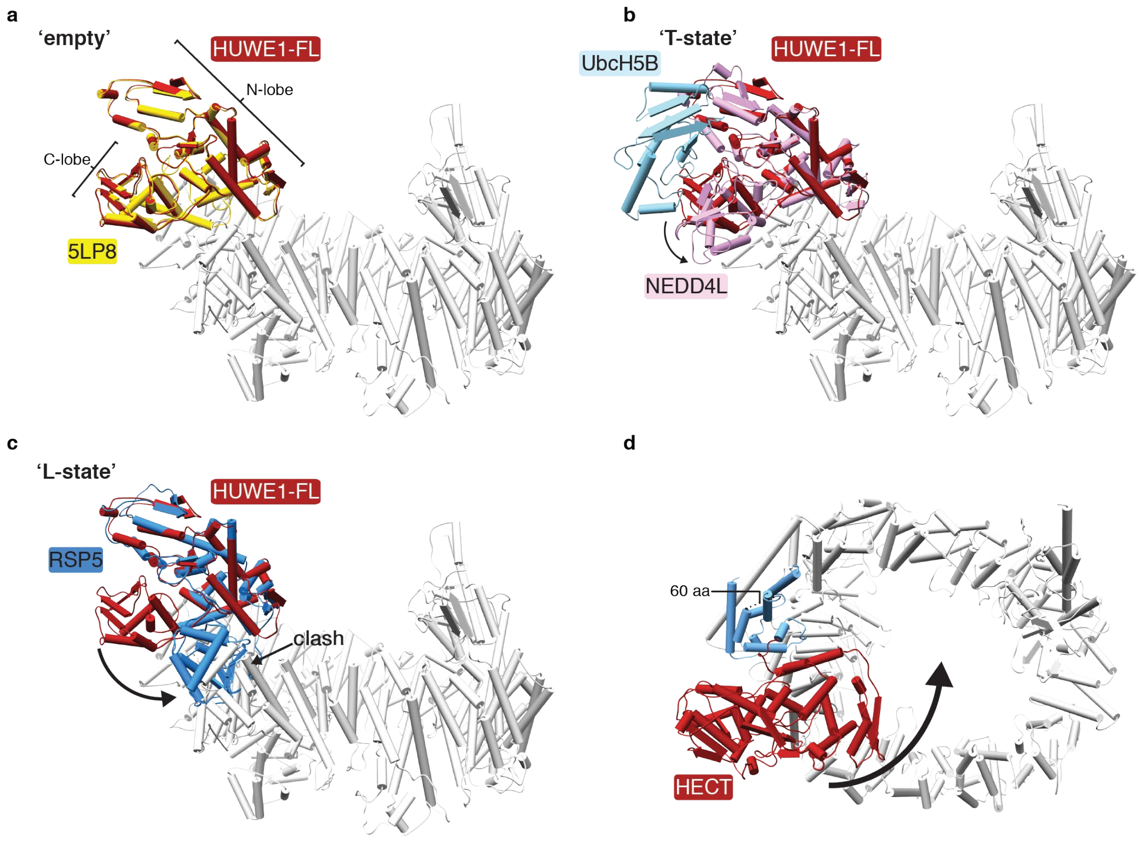

**Supplementary Fig. 6. HECT domain dynamics.** (**a**) Overlay of HUWE1 HECT (red) in context of full-length protein with HUWE1 HECT domain observed in isolation (yellow) in absence of binding partners[41,57] (PDB: 5LP8). (**b**) Overlay with a HECT/E2 complex (PDB: 3JVZ[58], HECT (NEDD4L): pink, E2 (UbcH5B): light blue). A small rotation (indicated by arrow) of the C-lobe would position the HUWE1 HECT domain (red) into a resting conformation commonly referred to as T-state for HECT ligases, which is hypothesized to allow for Ub-charged E2 binding. In this conformation, the active site cysteine, located in the C-lobe, is pointing away from the ring and all potential substrate binding domains. (**c**) Overlay with a transfer-trapped Rsp5-Sna3-Ubiquitin structure (PDB: 4LCD[59], only RSP5 is shown in dodger blue for better visibility). Transitioning HUWE1 HECT (red) domain into this conformation would lead to severe clashes with ARLD1. (**d**) Illustration of HUWE1 HECT domain (red) rotation (arrow) necessary to bring active site cysteine closer towards the center of the ring. Such a movement could be governed by rearrangements in C-terminal helices of ARLD4 (colored in blue) and is supported by structural variance analysis (**Supplementary Movie 1**).

**a**

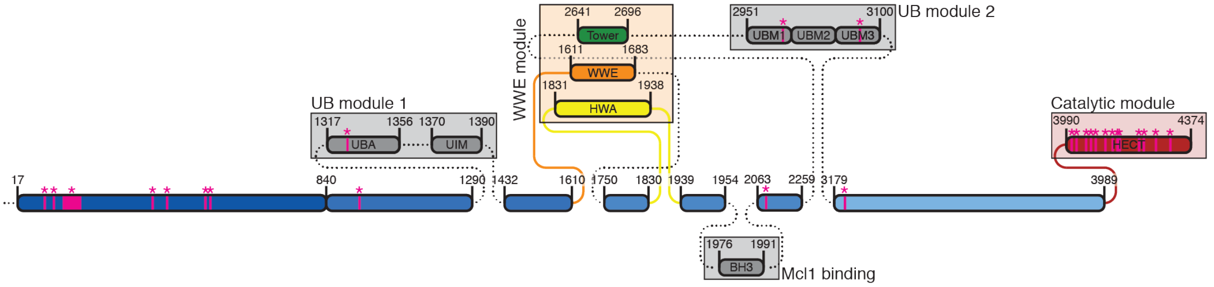

**b**

| | | |
|---|---|---|
| III | ARLD1 | R110W |
| III | ARLD1 | R110Q |
| III | ARLD1 | S115F |
| I | ARLD1 | del(169-189) |
| III | ARLD1 | A326G |
| III | ARLD1 | V433I |
| III | ARLD1 | G660R |
| III | ARLD1 | H669Q |
| III | ARLD2 | V950D |
| IV | UBA | M1328V |
| III | ARLD3 | R2040Q |
| III | ARLD3 | I2089M |
| IV | UBM | R2981H |
| IV | UBM | R3070C |
| I | ARLD4 | F3194S |
| II | HECT | R4013Q |
| II | HECT | R4023C |
| II | HECT | R4063Q |
| II | HECT | I4069F |
| II | HECT | N4075K |
| II | HECT | Y4106C |
| II | HECT | Y4157V |
| II | HECT | R4187C |
| II | HECT | R4237C |
| II | HECT | E4244D |
| II | HECT | K4295N |
| II | HECT | G4310R |

**c**

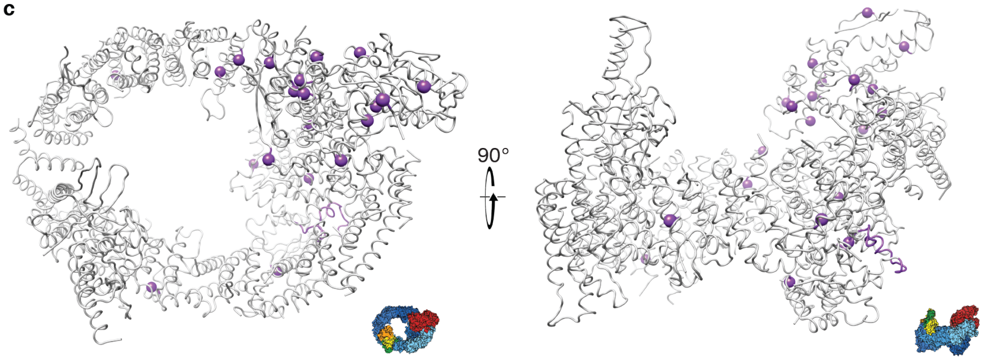

**Supplementary Fig. 7. Patient mutations in HUWE1.** (**a**) Schematic overview of HUWE1 with patient mutations described in Moortgat *et al*[3] labeled in pink. (**b**) Class, domain location and amino acid change of mutations. (**c**) Mutations mapped (purple spheres) onto HUWE1 structure.

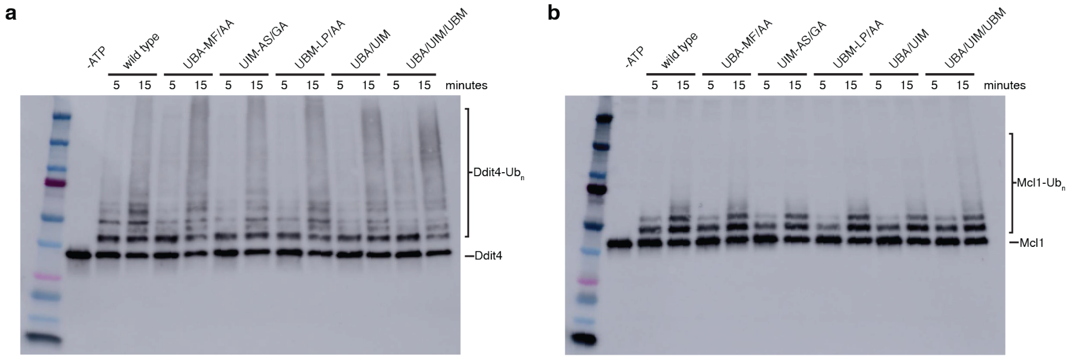

**Supplementary Fig. 8. Activity of UB module mutants.** (**a,b**) *In vitro* ubiquitylation of DDIT4 (a) or Mcl1 (b) by wild type HUWE1 or HUWE1 harboring mutations in the UBA (M1328A/F1330A)[60], UIM (A1376G/S1382A)[61], UBM (L4976A/P2977A)[62] domains, and combinations thereof. A slight increase in activity can be observed for mutants with combined mutations.

**Fig. 3**

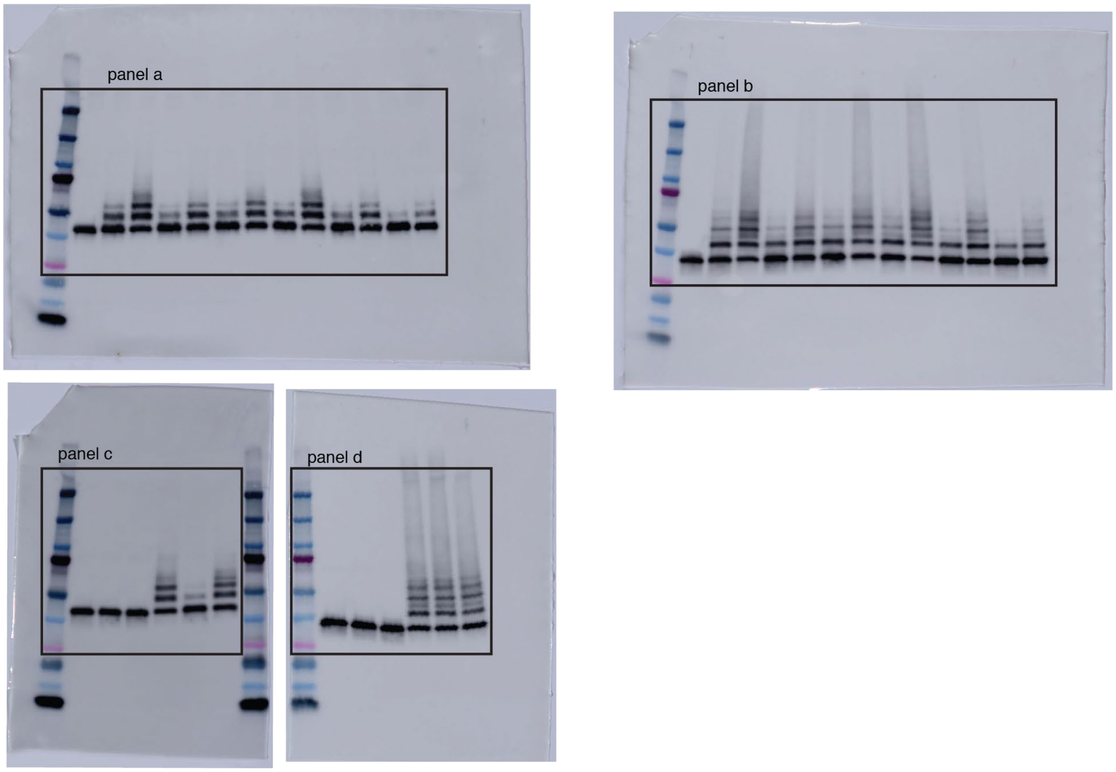

**Fig. 4**

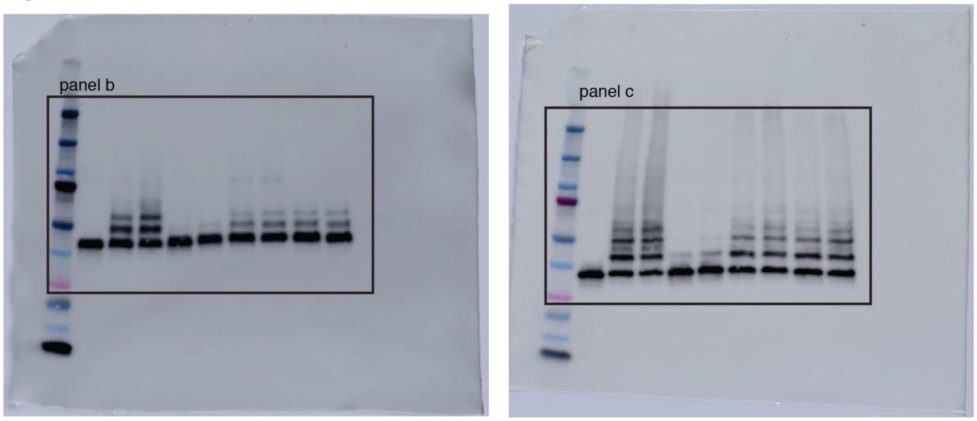

**Supplementary Fig. 9. Uncropped blots used in Figures 3 and 4.** Uncropped blots used in main figures with boxes indicating the respective area used in the figures.

**Supplementary Fig. 1**

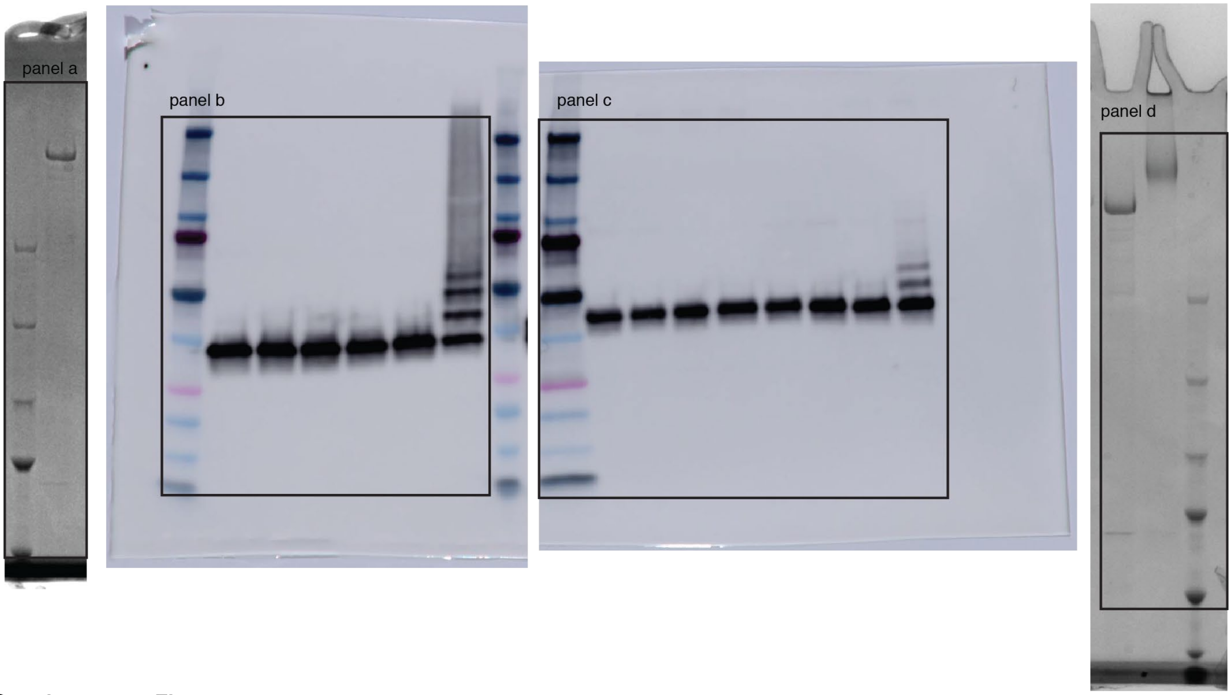

**Supplementary Fig. 7**

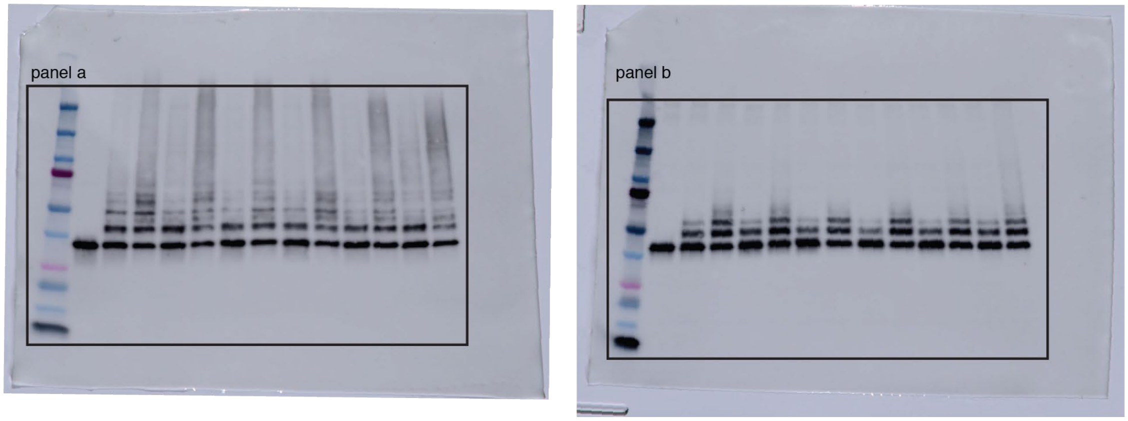

**Supplementary Fig. 10. Uncropped blots and SDS gels used in Supplementary Figures.** Uncropped blots used in Supplementary Materials Figures with boxes indicating the respective area used in the figures

#### Supplementary Table 1. Cryo-EM data collection, refinement and validation statistics

| | Data set 1<br>HUWE1-BS3<br>(EMD-22427)<br>(PDB: 7JQ9) | Data set 2<br>HUWE1 non-crosslinked<br>(EMD-22431) |
|---|---|---|
| **Data collection and processing** | | |
| Magnification | 105’000 | 130’000 |
| Voltage (kV) | 300 | 300 |
| Electron exposure (e–/Å$^2$) | 45.68 | 49.37 |
| Defocus range (μm) | -0.8 - -2.5 | -1 - -2.3 |
| Pixel size (Å) | 0.850* | 1.059† |
| Symmetry imposed | C1 | C1 |
| Initial particle images (no.) | 2’110’785 | 2’902’388 |
| Final particle images (no.) | 762’898 | 336’853 |
| Map resolution (Å)<br>FSC threshold | 3.1<br>0.143 | 3.4<br>0.143 |
| Map resolution range (Å) | 2.7 - 6 | 3 - 10 |
| **Refinement** | | |
| Initial model used (PDB code) | ab initio, 5LP8, 6MIW | - |
| Model resolution (Å)<br>FSC threshold | 3.2<br>0.5 | - |
| Map sharpening B-factor (Å$^2$) | -78.1 | - |
| Model composition<br>Non-hydrogen atoms<br>Protein residues<br>Ligands | <br>19165<br>2427<br>- | - |
| *B* factors (Å$^2$)<br>Protein<br>Ligand | <br>59.67<br>- | - |
| R.m.s. deviations<br>Bond lengths (Å)<br>Bond angles (°) | <br>0.007<br>0.935 | - |
| Validation<br>MolProbity score<br>Clashscore<br>Poor rotamers (%) | <br>1.66<br>6.82<br>1.68 | - |
| Ramachandran plot<br>Favored (%)<br>Allowed (%)<br>Disallowed (%) | <br>97.43<br>2.57<br>0 | - |

*later determined to be 0.825
†final pixel size after motion correction

**Supplementary Movie 1. 3D variability analysis of BS3-crosslinked HUWE1**

3D variability analysis (20 frames, filter resolution 5 Å, cryoSPARC v2.12.4) using the particles from the consensus refinement (EMD-22427), displaying HUWE1 from all sides. The HECT domain exhibits pronounced mobility, accompanied by a breathing motion of the entire alpha solenoid.

**Supplementary Movie 2. 3D variability analysis of the open form of non-crosslinked HUWE1**

3D variability analysis (20 frames, filter resolution 5 Å, cryoSPARC v2.12.4) using the particles marked in **Supplementary Fig 3a**, starting from the same view as in **Supplementary Movie 1**. The open form of HUWE1 is highly flexible and apparently in a continuous motion, prohibiting high-quality 3D reconstructions.

**References**

56 Tan, Y. Z. *et al.* Addressing preferred specimen orientation in single-particle cryo-EM through tilting. *Nat. Methods* **14**, 793-796 (2017).

57 Zhu, K. *et al.* Allosteric auto-inhibition and activation of the Nedd4 family E3 ligase Itch. *EMBO Rep.* **18**, 1618-1630 (2017).

58 Kamadurai, H. B. *et al.* Insights into ubiquitin transfer cascades from a structure of a UbcH5B approximately ubiquitin-HECT(NEDD4L) complex. *Mol. Cell* **36**, 1095-1102 (2009).

59 Kamadurai, H. B. *et al.* Mechanism of ubiquitin ligation and lysine prioritization by a HECT E3. *Elife* **2**, e00828 (2013).

60 Walinda, E. *et al.* Solution structure of the ubiquitin-associated (UBA) domain of human autophagy receptor NBR1 and its interaction with ubiquitin and polyubiquitin. *J. Biol. Chem.* **289**, 13890-13902 (2014).

61 Tanno, H. *et al.* Ubiquitin-interacting motifs confer full catalytic activity, but not ubiquitin chain substrate specificity, to deubiquitinating enzyme USP37. *J. Biol. Chem.* **289**, 2415-2423 (2014).

62 Bomar, M. G. *et al.* Unconventional ubiquitin recognition by the ubiquitin-binding motif within the Y family DNA polymerases iota and Rev1. *Mol. Cell* **37**, 408-417 (2010).
